# Sertraline enhances bacterial control by improving the pharmacodynamic - pharmacokinetic properties of frontline TB drugs

**DOI:** 10.1101/2025.07.13.664629

**Authors:** Kanika Bisht, Monica Yadav, Mahima Madan, Largee Biswas, Rajesh Pradhan, Rajeev Taliyan, Manjula Singh, Anita K. Verma, Vivek Rao

## Abstract

Given the need for innovative interventions for tackling the burden of TB, host directed therapies have emerged as a promising alternative in recent times. The combination of sertraline with frontline TB drugs has shown excellent promise in the murine models of infection in imparting better bacterial control and increasing host survival. We tested if the addition of sertraline worked to increase bacterial clearance in the random bred guinea pig model of TB infection that mimics the inter individual heterogeneity observed in the human response to infection. The combination of sertraline and frontline TB drugs effectively reduced bacterial burdens in the tissues of guinea pigs significantly better than the drugs alone with a marked betterment of lung histopathology. In order to evaluate the effect of sertraline on the pharmacodynamic properties of TB drugs, concentrations of the drugs were estimated in tissues at different time intervals over a period of 24h of administration to rats. Overall, addition of sertraline did not alter TB drug distribution or clearance from the animals, although enriching drug amounts transiently between 3-6 h in the different tissues. We thus highlight the advantage of an adjunct TB therapy with the inclusion of sertraline, an FDA approved antidepressant, in improving the PKPD of TB drugs and imparting better infection control in diverse models of infection.

## Introduction

Recent efforts towards TB control have focused on alternative pathogen-independent host-directed therapy regimens in order to effectively offset the development of drug resistance in the population [1-3]. New molecules with promise in bacterial control have been tested for their ability to control bacterial replication and prevent tissue damage in preclinical infection models like mice, guinea pigs, and non-human primates. Mouse strains allow for a detailed study of Mtb pathogenesis and the role of immune responses in infection control [4, 5]. However, mice have inherent disadvantages as test models for TB and therapy regimens: 1) being syngeneic, they do not portray the diversity of disease progression and infection response, and 2) are relatively resistant to disease with minimal resemblance to pathology seen in human TB [6]. The random-bred guinea pigs provide an optimal model for evaluating novel intervention strategies for TB essentially due to a high degree of susceptibility to infection, variety in response across animals, formation of lipid-rich necrotic granulomatous lesions similar to humans, and an excellent response to drug-mediated therapy [7-9].

One of the major problems is excessive toxicity associated with the prolonged TB therapy regimen, in part due to improper clearance and accumulation of the drugs in the host tissues [10]. It is therefore pertinent that any new drug moiety before its test in the human population is evaluated for its pharmacokinetics and pharmacodynamics (PK-PD) alone or in combination with frontline TB drugs. Further, the use of a combination of drugs as in the case of TB therapy, necessitates the evaluation of potential drug-drug interactions that could lead to toxic accumulation or aberrant removal of effective drugs from the host tissues, reducing drug effectiveness. Understanding the kinetic drug concentration in body tissues following the administration of a defined drug dose is thus vital for identifying effective drug doses without the development of drug resistance or toxicity, essential for the progress of new moieties in clinical trials.

We have previously demonstrated the promise of using sertraline (SRT) as an adjunct therapy for TB over the frontline TB drugs alone. The combination of sertraline with frontline TB drugs effectively reduces bacterial burden in the lungs of infected mice in chronic Mtb infection and also leads to enhanced host survival in an acute infection model [11]. Given the superior performance of a combination of anti-TB therapy (ATT) and SRT in controlling TB in murine models, we investigated the effectiveness of this combination in the more susceptible guinea pig model of infection. In addition, we assayed the kinetic distribution of the frontline TB drugs in the tissues of rats following administration of frontline TB drugs alone or in combination with sertraline at various intervals over a period of 24hrs. We demonstrate that SRT, akin to the murine model, effectively promotes bacterial clearance by the frontline TB drugs from guinea pig lungs and prevents excessive damage to the tissue. Moreover, the addition of SRT to the four frontline drugs promoted the initial PK-PD of the TB drugs, while the clearance rates were relatively unaltered in the different tissues of animals, demonstrating that the ATT-SRT adjunct combination regimen provides a superior yet safe regimen for faster TB control.

## Materials and Methods

### Reagents

The following reagents were procured from Sigma Aldrich, USA: Isoniazid (I3377), Pyrazinamide carboxamide (P7136), Ethambutol dihydrochloride (E4630), and Sertraline hydrochloride (S6319). Rifampicin (CMS18890) was procured from HIMEDIA laboratories, Mumbai, India.

### Guinea pig model of TB infection

All animal infections were conducted in an ABSL-3 facility as per the accepted recommendations of the IAEC (protocol no. 49/IAEC/VR&GK/Biochem/UDSC/28.11.2022). Specific pathogen-free guinea pigs were procured and housed in pathogen-free conditions after the requisite period of quarantine. Animals in equal proportion of males and females were infected with ∼100-1000 cfu of Mtb Erdman strain using an inhalation exposure system (Glas-Col, USA) for aerosol delivery of ∼ 100 cfu per animal. Treatment with antibiotics and SRT was initiated after 4 weeks of infection and provided *ad libitum* in the drinking water containing 2% sucrose twice a week. The animals were administered frontline TB antibiotics (INH-100 mg/kg, Rif-40 mg/kg, PZA-150 mg/kg, Eth-100 mg/kg), with or without (10 mg/kg) SRT. At specific time points, animals were euthanized, and tissues were aseptically harvested for bacterial CFU counting and histological examination. The lungs and spleens of the animals were lysed with sterile 1 μM glass beads in 10mL of saline in a Polytron homogenizer. The lysates were serially diluted, followed by plating on Middlebrook 7H11 agar plates, supplemented with 0.5% glycerol and 10% OADC to enumerate the bacterial loads. The accessory lung lobe was preserved for histological analysis and fixed in 10% formalin. Gross images of the 10% formalin-fixed lung lobes were captured using a Zeiss (Semi 2000®C) bright field microscope. Paraffin-embedded lung sections underwent H&E staining and scanned using an Olympus microscope.

Whole animal (oral dosing) studies were done to determine the dose response curves or predict non-linear dose response curves. Samples were collected from the tissues at 0.5, 1, 2, 4, 6, and 24 h after administration of a single dose of the drugs.

### Analysis of PKPD

Wistar Rats were used in a certified animal facility as per the accepted recommendations of the IAEC (protocol no. DU/KR/IAEC/2023/16) for the pharmacokinetic studies. For analysis of PK-PD of TB drugs, a single administration of oral dose was conducted in 36 Wistar rats divided into six dosing groups. Briefly, 3 Wistar rats were given a single oral dose of a combination of drugs INH (100 mg/kg), Rif (40 mg/kg), Eth (100 mg/kg), PZA (150 mg/kg) and SRT (3 mg/kg) in a total volume of 1 mL. Rats were sacrificed at 3 different time points, i.e., 2 h, 4 h, and 24 h to evaluate the biodistribution of drugs in blood (serum) and tissues (Kidney, Lung, Liver, and Brain) and were collected in tubes containing PBS. Samples were analyzed for the drug concentrations by LC-MS in a triple quadrupole mass spectrometer, equipped with an electrospray ionization probe LC-MS/ MS system controlled by MassLynx V4.2 software (SCN1019). The concentrations were calculated by using a standard curve. For the standard curve, the compounds were dissolved at the required concentration in MS-grade water to generate a standard curve and injected (10 μL) into the Waters Acquity UPLC BEH C18 column (1.7 μm, 2 x 100 mm) at a flow rate of 0.450 mL/min at 35 °C with a total run time of 5.2 mins. The mobile phase was a gradient of solvent A (10 mM ammonium formate in water) and solvent B (0.1% formic acid in acetonitrile) along with a column cleaning gradient (table 1). The non-compartment model was used to calculate the pharmacokinetic parameters with PKSolver. The parameters used to characterize the disposition of a test substance were half-lives of elimination and absorption; area under the concentration-versus-time curve (AUC) for blood; total body; volume of distribution (Vd); and mean residence time (MRT). For the oral route, concentration at time zero was assumed to be zero. The plasma and tissue concentrations below the limit of quantitation were treated as null samples for the purpose of calculating the mean plasma concentration values or for calculating PK parameters. The area under the curve (AUC) versus time was calculated using the linear trapezoidal method (linear interpolation).

## Results

### Sertraline enhances bacterial clearance from the infected host

We have previously demonstrated that the addition of sertraline in the Mtb-murine infection model facilitates early bacterial clearance by frontline TB drugs. To test whether sertraline affects Mtb control in the random-bred guinea pig infection model, animals infected with 500 CFU were treated with the frontline antibiotics HRZE, with and without sertraline, and the bacterial numbers were enumerated at 5W and 10W post-treatment. A combination of sertraline and HRZE imparted significantly better bacterial control as early as 5 weeks (Figure 1A). Lungs of the animals treated only with antibiotics showed marginal restriction of bacterial numbers (0.5 logs) by 5 weeks of treatment as opposed to the > 10-fold decrease in the animals treated with HRZE along with sertraline. Importantly, most of the animals (3/4) harbored lower bacteria in the spleens of the HRZES-treated group, in comparison to the HRZE-treated animals. While at 10 weeks of treatment, the antibiotics alone showed ∼10-fold control and the addition of sertraline did not alter the bacterial numbers significantly in the lungs, better bacterial control could be observed in the spleens of sertraline treated animals (Figure 1B).

**Figure 1:**
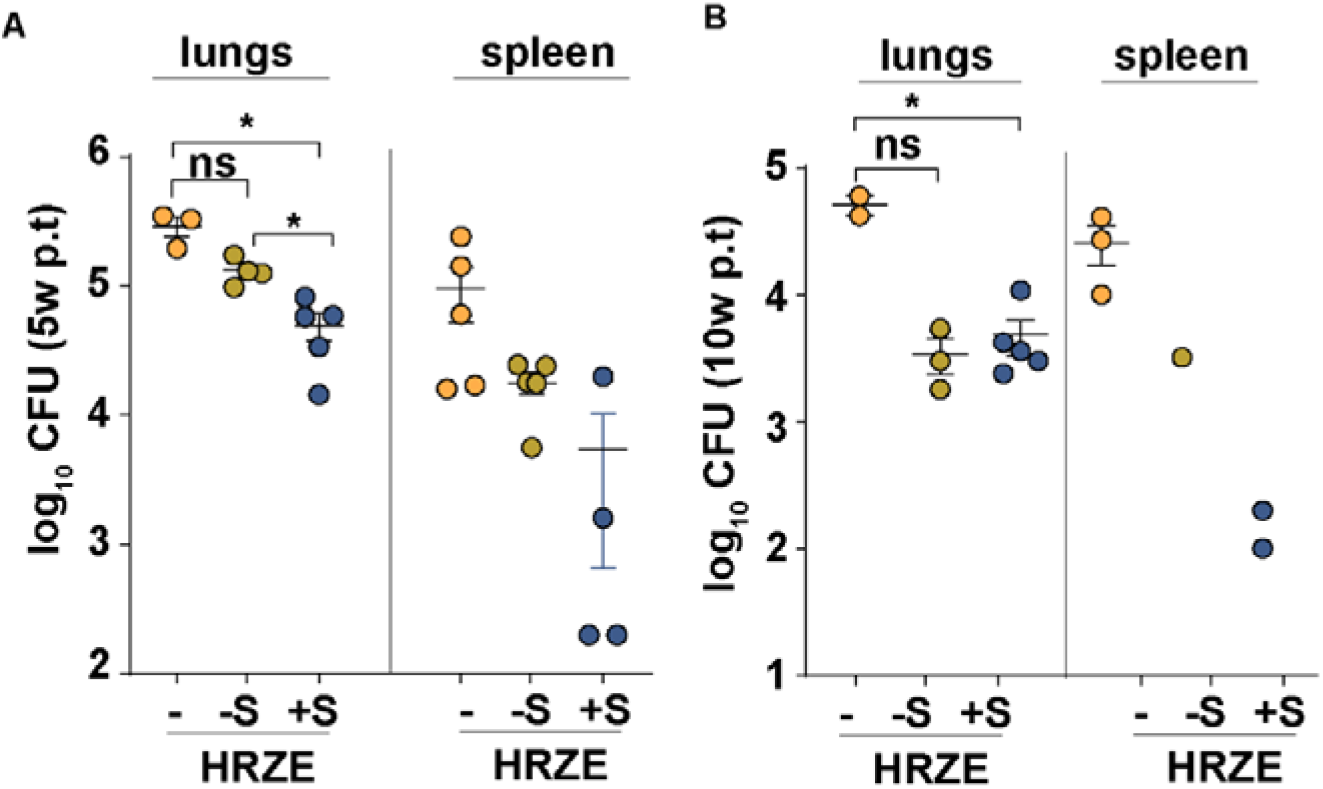
Effect of sertraline adjunct regimen in the guinea pig model of Mtb infection. The bacterial numbers in lungs and spleens of Mtb-infected, HRZE or HRZES-treated animals were enumerated by plating homogenates at 5w (A) and 10w (B) post-treatment. The values are mean values of CFU/ tissue ± SEM from 5 individual animals (N=5).

### Addition of sertraline to ATT prevents excessive tissue damage associated with infection

Infection with Mtb results in granuloma formation, the characteristic accumulation of immune cells in the tissues [12, 13]. While this response is aimed at infection spread and consequent control, uncontrolled involvement of the lung results in extensive tissue consolidation, resulting in respiratory distress [14, 15]. In line with our previous observation with the murine model, inclusion of sertraline as a combination regimen with frontline TB drugs significantly impacted lung histopathology of infected guinea pigs (Figure 2) as evident in the gross morphology of the lung tissues wherein the extent of macroscopic lesions was significantly higher for the untreated (NT) animals at both 5w and 10w time points with a slight betterment upon treatment with HRZE (Figure 2A). In contrast, most animal lungs showed evidence of lesser involvement of granulomatous response with morphologically decreased lesions in the HRZES-treated animals. This betterment of lung tissue was mostly evident in the histological sections of the lungs from the animals (Figure 2B). Again, while the tissue sections of both NT and HRZE groups of animals harbored significantly higher numbers of microscopic lesions that increased in numbers from 5w to 10w, significantly lower numbers of lesions were observed in the lung sections of HRZES-treated animals at both the time points, again highlighting the enhanced effect of adding sertraline as an adjunct to the TB therapy regimen for enhanced bacterial control and tissue recovery.

**Figure 2:**
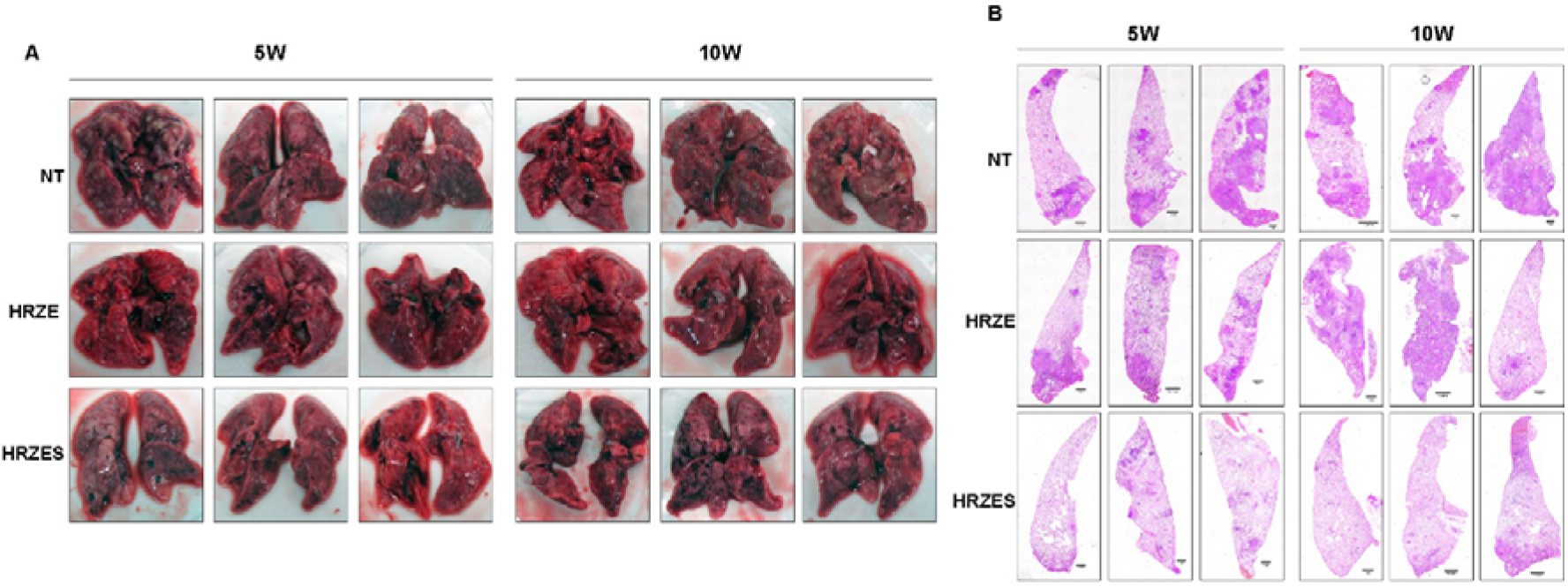
Sertraline protects extensive tissue damage in lungs. The gross morphology of lungs is depicted in the images after 5w and 10w of treatment (A). The lungs of animals were subjected to histopathology and H&E staining for evaluation of the microscopic lesions and consolidation of the tissue (B).

### Establishment of an assay system for dynamic analysis of frontline TB drug concentrations

Given the success of sertraline as an adjunct for standard TB therapy in augmenting bacterial control as well as preventing tissue damage associated with Mtb infection in preclinical models, we sought to check if the inclusion of sertraline altered the pharmacokinetic-properties of the frontline TB drugs. For this, the biodistribution as well as the effective drug concentrations of the TB drugs were evaluated at different intervals over a period of 24 h in the serum and tissues of rats treated with HRZE or HRZES. In order to establish the bioassay for quantitation of the drugs and sertraline, initial steps involved the generation of a standard calibration curve for the purified individual drugs. The calibration curves for all the analytes were best fitted with a quadratic regression weighted 1/c^2^. A single linear curve was established to facilitate drug quantification with 6 points (from 1 ng/mL to 200 ng/mL (approx. 5-200 ppm)) for all the drugs used in the study (Figure S1), The equations derived for estimation of tissue concentrations of the drugs is depicted in table 2.

**Table 2:** The standard curve equation and correlation coefficients of Rifampicin, Pyrazinamide, Isoniazid, Ethambutol and Sertraline.

| Analytes | Standard curve Equation | R<sup>2</sup> value |
| --- | --- | --- |
| Rifampicin | 3101.24x +-7751.26 | 0.9955 |
| Pyrazinamide | 73.035x + 941.59 | 0.9936 |
| Isoniazid | 1154.67x +2787.72 | 0.9984 |
| Ethambutol | 10375x + -1522.99 | 0.9978 |
| Sertraline | 11041x + -22028.4 | 0.9958 |

### Sertraline does not affect the long-term clearance or the kinetic distribution of frontline TB drugs in serum

A combination of molecules is often reported to affect the accumulation and clearance of the individual compounds from the host. To test if sertraline modifies long-term clearance of frontline TB drugs, their concentrations were estimated in animals treated for 24 h with the drugs alone or in combination with sertraline at 2 doses of 3mg/kg and 1mg/kg. As observed in Figure 3, despite the absence of any detectable levels of rifampicin across treatments, the addition of sertraline did not alter the levels of frontline TB drugs in animal sera. The concentrations of isoniazid and pyrazinamide were similar between the HRZE, HRZES1, and HRZES2 groups of animals at both the initial 30 min and later 24 h time points after treatment. While ethambutol concentrations were initially higher (not significant) in the HRZES2 at 0.5 hrs after administration, the levels returned to more or less normal levels. Given the similar levels of the drugs at 24 hrs, we asked if the combination with sertraline modified the temporal availability of drugs in the serum by evaluating the drug concentrations at 1h, 2h, 4h, and 6h after drug administration in the animals. Again, at either concentration, the presence of sertraline did not significantly change the levels of the TB drugs at any time point (Figure 3). Taken together, these data highlight the insignificant impact of sertraline on serum concentrations of frontline TB drugs.

**Figure 3:**
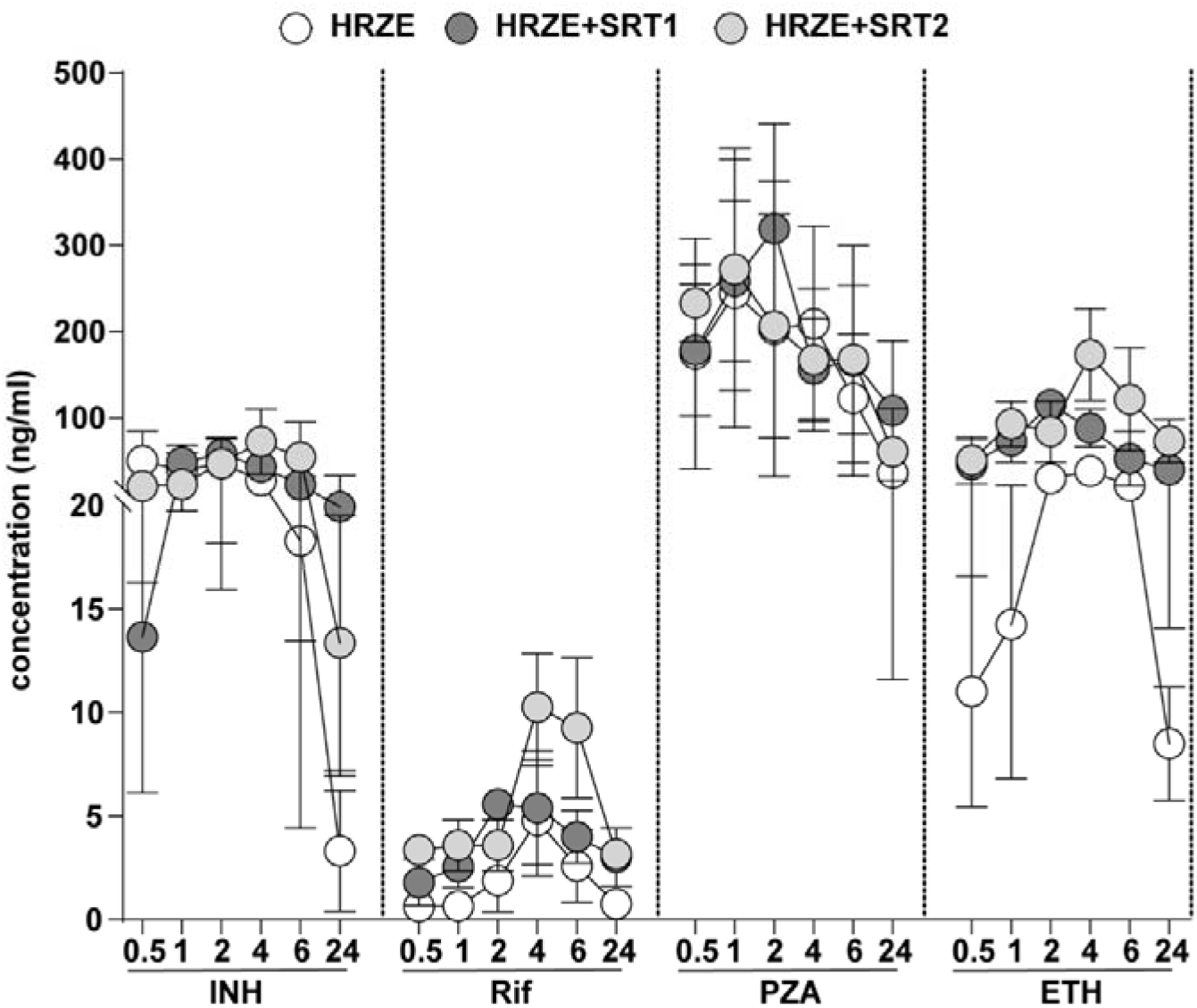
Distribution of frontline TB drugs in animals. Estimation of concentrations of Isoniazid (INH), Rifampicin (Rif), Pyrazinamide (PZA) and Ethambutol (ETH) in serum samples of rats after oral administration of the drugs alone (HRZE) or in combination with sertraline at two different concentrations of 3mg/kg (HRZES1) and 1mg/kg (HRZES2). Rats were bled at 30’ and at various intervals after treatment, and the compounds were estimated in the sera by LC-MS. Values represent the mean ± SEM of the relative concentrations (rel. to T_30’_) of 3 independent animals (N=3).

### Negligible impact of sertraline on the distribution and retention of TB drugs in the lungs of treated animals

Sertraline has been shown to effectively enhance the antimicrobial properties in pulmonary TB in murine model infection [11]. Given the effect of sertraline on lung infection, we questioned if distribution of TB drugs in this tissue was altered by the combination therapy. Surprisingly, the accumulation of TB drugs was not altered by sertraline at most of the intervals studied (Figure 4). Interestingly, the level of PZA was significantly higher at the most time points in the sertraline treated group as compared to the drug alone treated animals.The clearance of the drugs was completely unaffected by the presence of sertraline in the lungs of treated animals.

**Figure 4:**
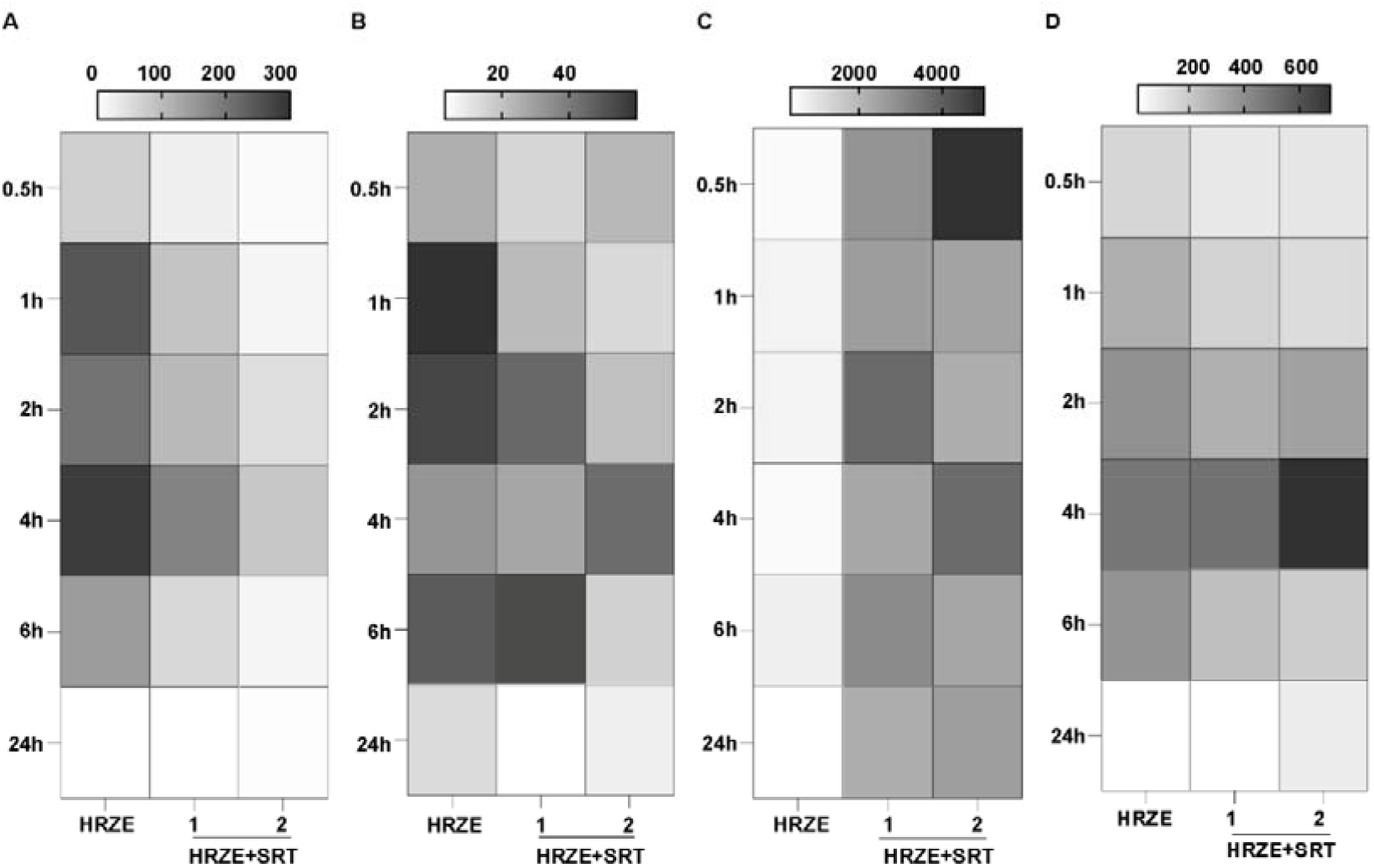
Kinetic profile of lung concentrations of frontline drugs in vivo. The relative concentrations in the lungs of treated rats of isoniazid (INH, A), rifampicin (Rif, B), pyrazinamide (PZA, C), and Ethambutol (ETH, D) were estimated by LC-MS at intervals of 0.5 h, 1 h, 2 h, 4 h, and 6 h post-drug administration. Individual drug levels in animals receiving the TB drugs (HRZE) were compared with animals receiving a combination of HRZE with 3mg/kg (HRZES1) and 1mg/kg (HRZES2) of sertraline. Values represent the mean **±** SEM of the relative concentrations (rel. to T_30’_) of 3 independent animals (N=3).

### Addition of sertraline does not disturb the temporal distribution or clearance of frontline TB drugs

New chemical entities like drugs introduced into the mammalian systems are eliminated through detoxification by the liver and excretion by the kidneys, failing which drugs accumulate in the system, leading to significant associated toxicities. To test if the presence of sertraline affects drug removal, the levels of the drugs were estimated in the livers and kidneys of treated animals. Again, the absence of a detrimental effect of sertraline on the TB drug clearance was evident in the animals. All the frontline TB drugs reached comparable levels by 24 h of introduction (Figure 5) in the liver and kidneys of the animals. In fact, the levels of all the four drugs were comparable in both the control (HRZE) group and the combination groups with sertraline (HRZES1 and HRZES2) at all time points in the liver (Figure 5A) and the kidneys (Figure 5B) despite the relatively short lined inflection in pyrazinamide levels at 0.5 hrs. Sertraline is an FDA-approved antidepressant with efficient capability to penetrate the blood-brain barrier (BBB), thereby enabling it to control CNS infections either alone or in combination with antibiotics. To verify if sertraline induced the accumulation of drugs in the brains of treated animals, the concentrations of the drugs in the brain were determined periodically until 24 h. As observed in the case of other tissues, addition of sertraline in either concentration did not result in the excess accumulation of the drug in the brain (Figure 5C). The drug concentrations were comparable in the groups treated with HRZE alone or in combination with sertraline at the initial 30’ as well as the late 24 h time point. Cumulatively, these results effectively provide evidence for the absence of any deleterious effect of sertraline addition on the accumulation and clearance of frontline TB drugs in vivo, thereby highlighting the safety of the combination therapy.

**Figure 5:**
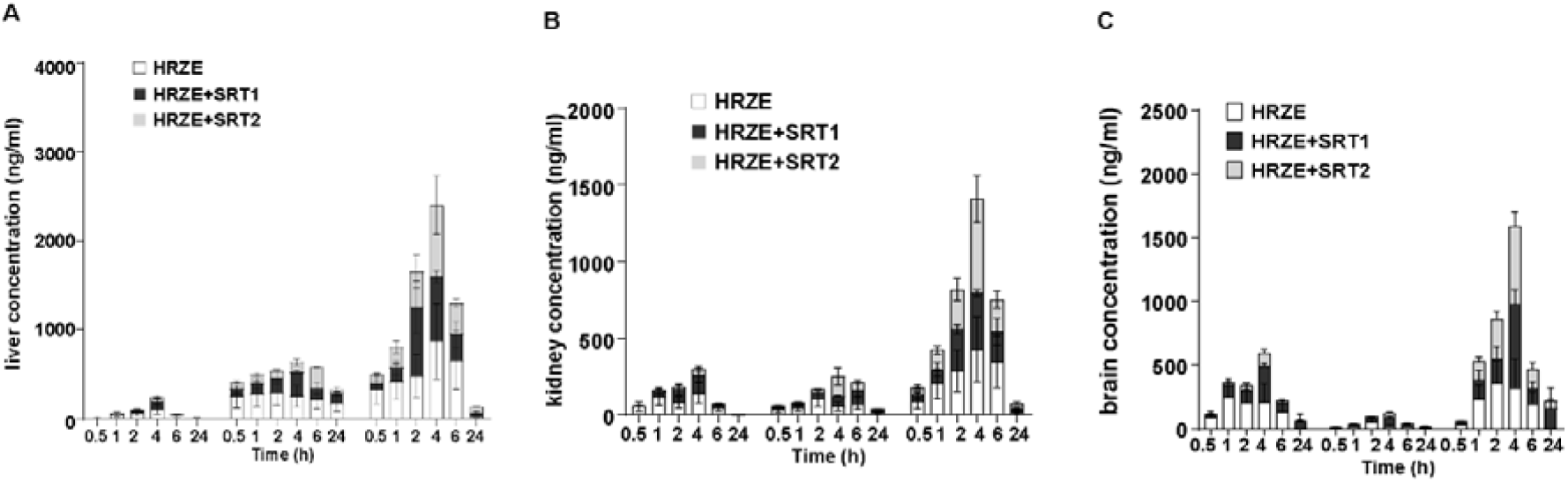
Tissue concentrations of frontline TB drugs. The relative concentrations of drugs in the liver (A), kidneys (B), and brain (C) of treated rats were estimated by LC-MS at intervals up to 24 h as indicated post-drug administration. The levels in animals receiving the TB drugs (HRZE) were compared with animals receiving a combination of HRZE with 3mg/kg (HRZES1) and 1mg/kg (HRZES2) of sertraline. Values represent the mean ± SEM of the relative concentrations (rel. to T_30’_) of 3 independent animals (N=3).

## Discussion

The recent trend of harnessing host mechanisms for effective control of infectious diseases (host-directed therapies, HDTs) has seen the advent of multiple novel entities as putative drugs [1, 16, 17]. More so, given the long duration of the current TB regimen and the associated drug resistance, HDTs offer an excellent alternative that can circumvent this problem. In fact, numerous modalities have been devised as effective HDTs for TB in recent years [18-20]. Previously, we have demonstrated the effectiveness of adding the FDA approved antidepressant, sertraline, to standard TB drugs in achieving faster control of bacterial growth [11]. In the preclinical murine model, a combination of sertraline with the frontline TB drugs resulted in early clearance of Mtb from the host in addition to a significant improvement of the tissue from infection-induced injury (granuloma). While the syngeneic strains of mice are readily infected by Mtb and mirror the immune responses generated in humans, the inherent lack of variability and differences in the disease progression with that seen in human TB necessitate the testing of newer intervention strategies in a more disease-relevant preclinical model. The drastic increment in survival of Mtb-infected mice prompted the testing of this adjunct regimen in the guinea pig model of aerosol TB infection. In line with our observations with the murine infections, we observed a significant enhancement of bacterial control by the addition of SRT to the standard TB drugs, which reflected as a marked reduction in granuloma and an improved lung architecture.

It is not surprising that the majority of individuals infected with Mtb fail to manifest the disease, while some individuals portray varying degrees of lung consolidation and granulomatous lesions [21, 22]. This was also evident in the outbred and genetically diverse guinea pig model of infection, with lungs of infected animals displaying different degrees of consolidation with granulomatous lesions [7, 23]. Despite this associated heterogeneity in response kinetics, the response of animals in clearing the infection with the combination was significantly similar, with almost all the animals treated with HRZES showing lesser bacterial numbers than the antibiotic alone (HRZE) group, and most animal lungs showing clear, homogenous lungs with the characteristic alveolar spaces. This provides a superior basis for the possible effectiveness of the combination in a more genetically and phenotypically diverse human population.

Sertraline has been tested as an adjunct therapy for several infections and has shown promise in enhancing control of bacterial as well as viral infections [24-27]. In fact, by virtue of its ability to cross the BBB, it has been recognized as a promising candidate for the intervention of infections of the CNS. Recent studies have demonstrated the ability of SRT to significantly advance the control of neurotropic *Cryptococcal* infection in murine brain [28, 29].

Given its ability to enhance ATT efficacy, it is essential to evaluate the safety of the combination prior to testing in the human population.

While animals given the combination recovered faster from infection with no obvious indications of toxicity, direct evaluation of possible deleterious effects of sertraline on TB drug bio-distribution or clearance, resulting in toxic buildup of the drugs, is imperative. To investigate this, we tested the dynamic concentrations of the TB drugs in rats across 24 h of TB-drug administration either with or without sertraline. At different time intervals, the concentration of the TB drugs in serum, lungs, brain, kidneys, and liver was estimated by LC-MS.

Most of the TB drugs have associated toxicity with long-term usage, particularly isoniazid shows a usage-dependent liver toxicity [30]. One of the effective methods to offset this toxicity is a reduction in treatment durations, which can be achieved by newer modalities with faster killing kinetics or enhancing the bioavailability of the drugs without affecting the clearance of drugs from the system. Free serum rifampicin distributes well throughout body tissues, but concentrations in various tissues vary, with concentrations often lower than those in blood [31]. Our results indicate that the concentration of each drug differed greatly in each organ. With the addition of sertraline, higher level of INH was observed in the liver, lungs, brain, and kidneys up to 6 h, which was eliminated after 24 h, thereby satisfying the latter condition of absence of accumulation and providing a promising alteration in drug concentration with combination therapy. Even for ethambutol, the concentrations in the liver, kidneys, lungs and brain showed a partial increase after 4 h in the presence of SRT1 and SRT2, with complete elimination from the tissues by 24 hrs. A similar profile was also observed for rifampicin, one of the most potent TB drugs, with a comparable increase in the tissues, after 4 h in the presence of SRT1 and SRT2, except for the kidneys, where a higher concentration was observed after 4 h only with SRT2. Again, all the drugs were eliminated from the animal tissues by 24 h highlighting the improvement of the PK-PD of TB drugs with the addition of SRT.

In contrast, the concentration of pyrazinamide in the tissues showed a partial increase in the presence of SRT1 and SRT2 in comparison to the absence of SRT from 0.5 h to 24 h. Interestingly, PZA shows persistence even after 24 h in all tissues, without the addition of sertraline, indicating a slower kinetics of removal of the drug from the host. In conclusion, there was an increase in PZA in both kidneys and brain in the presence of SRT1 and SRT2. PK analysis indicated that PZA showed persistence at 24 h, probably due to the higher dose given (150 mg/kg). There was an increase in AUC and MRT of PZA in the presence of SRT1 and SRT2. However, the T_max_ of all the TB drugs never increased beyond 4h of administration with SRT, suggesting that the slight enhancement of the concentration with SRT and the decline over time would be insignificant for any hazardous buildup of PZA in the animals (data not shown).

In conclusion, our results suggest a positive impact of adding sertraline to TB drugs with a transient increase in bioavailability in the serum as well as in the tissues without any additional toxic accumulation of the TB drugs in the host over and above the levels of the 4-drug combination alone. We also observed a significant increase in the antibiotic control of Mtb in the guinea pig model of infection with the addition of sertraline. While it is plausible to assume that the initial increase in the antibiotic concentrations in the serum as well as in the tissues would result in the increased benefit observed with SRT in bacterial control, more detailed studies would provide this linear relationship in appropriate models of infection. Overall, we have developed a simple, sensitive, and cost-effective LC-MS/MS method for the simultaneous quantification of sertraline and other tuberculosis drugs and provided initial insights into the mechanisms of enhancing TB treatment in an attempt to reduce the duration of therapy.

The calibration ranges of the developed method covered the proposed therapeutic concentration ranges for the analytes.

**Figure S1:**
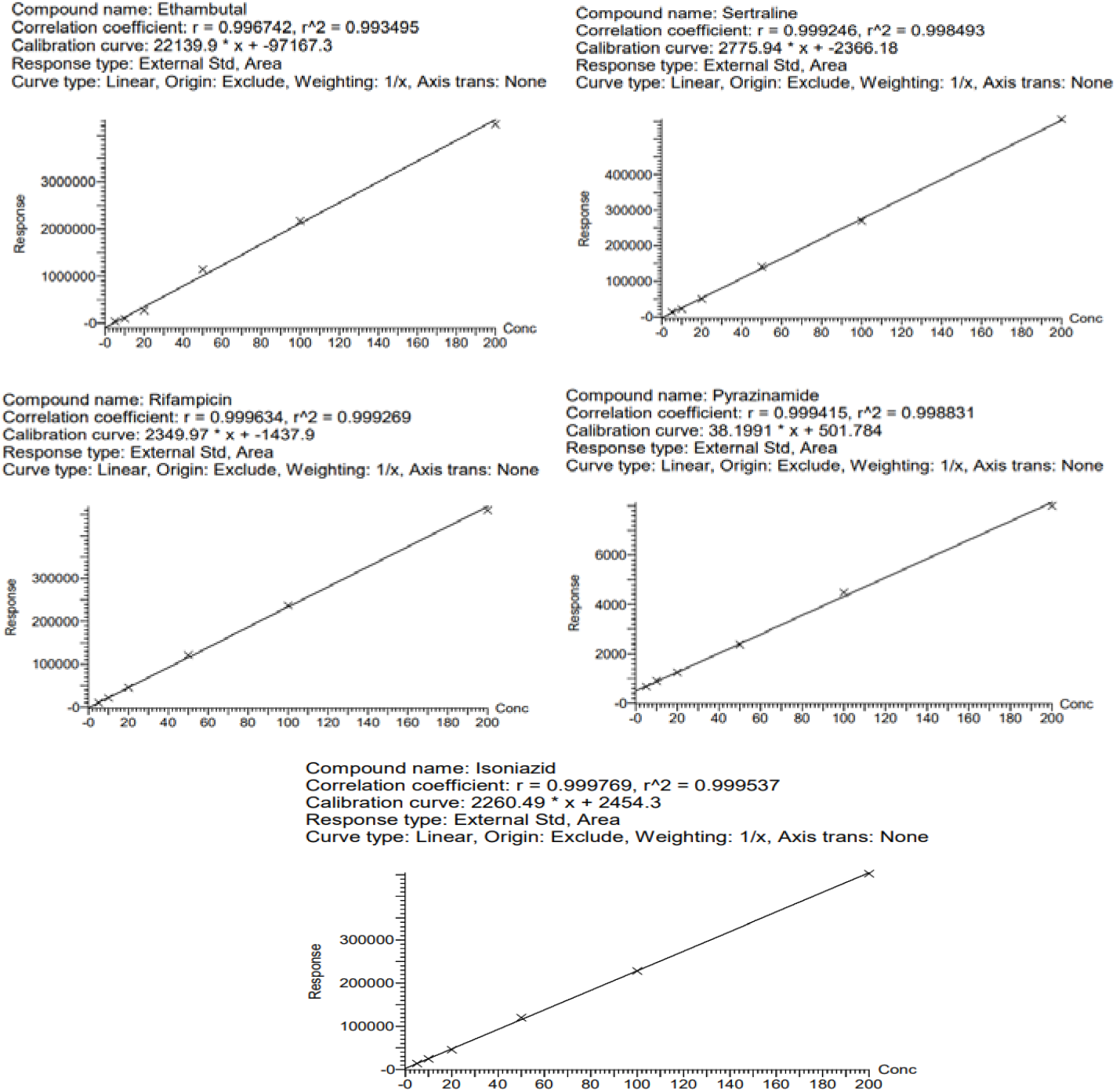
Standardization/Optimization of assays for detection of the frontline TB drugs and sertraline by LC-MS.

## Ethics

Animal work-the work was carried as per the requisites of the institutional animal ethics committee, approval (IGIB/IAEC/13/28/2020 and 49/IAEC/VR&GK/BIOCHEM/UDSC/ 28.11.2020).

## Author contribution

KB, LB, RP, RT, MS, AV and VR were instrumental in the design of the work. KB, MY, MM, LB, RP were involved in the conduct of the work. None of the authors have any conflict of interest.

## Acknowledgements

The authors thank ICMR (VR-GAP0213) for supporting the study. The student fellowships from CSIR-India are acknowledged.

